# Insect herbivory on urban trees: Complementary effects of tree neighbours and predation

**DOI:** 10.1101/2020.04.15.042317

**Authors:** Alex Stemmelen, Alain Paquette, Marie-Lise Benot, Yasmine Kadiri, Hervé Jactel, Bastien Castagneyrol

**Affiliations:** BIOGECO, INRAE, Univ. Bordeaux – 33610 Cestas, France; Département des sciences biologiques, Centre d’étude de la forêt (CEF), Université du Québec à Montréal – Centre-Ville Station, P.O. Box 8888, Montréal, Qc H3C 3P8, Canada

**Keywords:** Artificial prey, Insect herbivory, Tree diversity, Top-down control, Urban biodiversity

## Abstract

1. Insect herbivory is an important component of forest ecosystems functioning and can affect tree growth and survival. Tree diversity is known to influence insect herbivory in natural forest, with most studies reporting a decrease in herbivory with increasing tree diversity. Urban ecosystems, on the other hand, differ in many ways from the forest ecosystem and the drivers of insect herbivory in cities are still debated.
2. We monitored 48 urban trees from five species – three native and two exotic – in three parks of Montreal (Canada) for leaf insect herbivory and predator activity on artificial larvae, and linked herbivory with both predation and tree diversity in the vicinity of focal trees.
3. Leaf insect herbivory decreased with increasing tree diversity and with increasing predator attack rate.
4. Our findings indicate that tree diversity is a key determinant of multitrophic interactions between trees, herbivores and predators in urban environments and that managing tree diversity could contribute to pest control in cities.

This article has been peer-reviewed and recommended by *Peer Community in Ecology* https://doi.org/10.24072/pci.ecology.100061

## Introduction

Insect herbivores have a major impact on tree growth and survival, hence on the functioning of forest ecosystems (Metcalfe et al., 2014; Visakorpi et al., 2018; Zvereva, Zverev, & Kozlov, 2012). Tree diversity significantly influences insect herbivory in forest ecosystems (Castagneyrol, Jactel, Vacher, Brockerhoff, & Koricheva, 2014; Jactel et al., 2017). Most studies report that herbivory declines as tree diversity increases (*i.e*., associational resistance, Barbosa et al., 2009), although the opposite pattern has also been found (Haase et al., 2015; Schuldt et al., 2011). Recently, the interest in how tree diversity affects insect herbivory has expanded to include urban forests (Clem & Held, 2018; Dale & Frank, 2018; Frank, 2014), where pest damage can compromise the ecological and aesthetic values of urban trees (Nuckols & Connor, 1995; Tooker & Hanks, 2000; Tubby & Webber, 2010). Urban forests differ from natural forests in many ways. For example, most of the trees in cities are planted, found in lower density and/or mixed with native and exotic ornamental species that are rarely encountered in natural forests. Thus, given these specific characteristics of urban forests, it is still unclear how and why tree diversity might influence insect herbivory on urban trees.

The density and diversity of trees determine the amount and the quality of food and habitat resources available to herbivores and their enemies, and thus can have strong impact on the bottom-up and top-down forces acting upon insect herbivores (Haase et al., 2015; Muiruri, Rainio, & Koricheva, 2016; Setiawan, Vanhellemont, Baeten, Dillen, & Verheyen, 2014). For example, some insect herbivores, in particular generalist species, could take advantage of tree diversity to acquire more abundant, complementary food resources or benefit from a more balanced food mix, thus causing more damage in mixed forests (Lefcheck, Whalen, Davenport, Stone, & Duffy, 2013). In contrast, insect herbivores generally find it easier to identify and orientate towards the signals emitted by their host trees when the latter are more concentrated (*the resource concentration hypothesis*, Hambäck & Englund, 2005; Root, 1973) while non-host trees can emit volatile compounds that interfere with the ability of herbivores to detect their preferred host (Jactel, Birgersson, Andersson, & Schlyter, 2011). Finally, the abundance and diversity of predatory birds and arthropods generally increases with plant density and diversity, which would result in a better top-down regulation of insect herbivores (*the enemies hypothesis*, Risch, Andow, & Altieri, 1983; Root, 1973). However, the evidence available to support the enemies hypothesis in forest is controversial (Muiruri et al., 2016; Riihimäki, Kaitaniemi, Koricheva, & Vehviläinen, 2005; Staab & Schuldt, 2020) and the contribution of natural enemies to the control of herbivores in urban area remains poorly explored.

Tree diversity and density vary widely between and within cities (Ortega-Álvarez, Rodríguez-Correa, & MacGregor-Fors, 2011; Sjöman, Östberg, & Bühler, 2012). A consequence of this variability is that even within a common urban environment, herbivory may be reduced in some tree species and increased in others (Clem & Held, 2018; Frank, 2014), and the relative importance of bottom-up and top-down forces responsible for these effects may also differ. In addition, non-native trees have been widely planted in urban habitats (Cowett & Bassuk, 2014; Moro, Westerkamp, & de Araújo, 2014). While they often escape from herbivory by native insects (‘*the enemy escape hypothesis*’, Adams et al., 2009; Keane & Crawley, 2002), cases of native herbivores spilling-over onto exotic trees have been recorded (e.g. Branco, Brockerhoff, Castagneyrol, Orazio, & Jactel, 2015). Non-native tree species can also provide habitats to insectivorous birds or predatory arthropods (Gray & van Heezik, 2016). It is thus difficult to predict the effect of mixing native and exotic trees on insect herbivory in urban habitats (Clem & Held, 2018; Frank, 2014).

In this study, we investigated the effect of tree density, tree diversity, presence of conspecific trees, tree origin and predator activity on insect herbivory in urban trees of the city of Montreal (Quebec, Canada). We measured leaf area removed or otherwise damaged by insect herbivores on 48 trees of five species – three native and two exotic – in three urban parks. We concomitantly assessed predator activity by using artificial caterpillars exposed on tree branches. We tested the following hypotheses: (1) insect herbivory decreases with tree density, number of non-conspecific trees (host dilution) and diversity (associational resistance) around focal trees, (2) predator activity increases with increasing tree density and diversity and (3) predation and herbivory have different responses to tree diversity on native and exotic trees. By doing so, our study builds toward a better understanding of the drivers of pest insect damage on urban trees.

## Methods

### Study site

The study was conducted in the city of Montreal (Canada, 45°50’N, -73°55’W), where the climate is temperate cold, with 6.8°C average temperature and 1000.3 mm annual rainfall during the 1981-2010 period (Pierre Elliott Trudeau airport weather station, www.canada.ca). The study took place in three parks of the southwest part of the city: Angrignon, Marguerite Bourgeoys and Ignace-Bourget (Table 1).

**Table 1.**
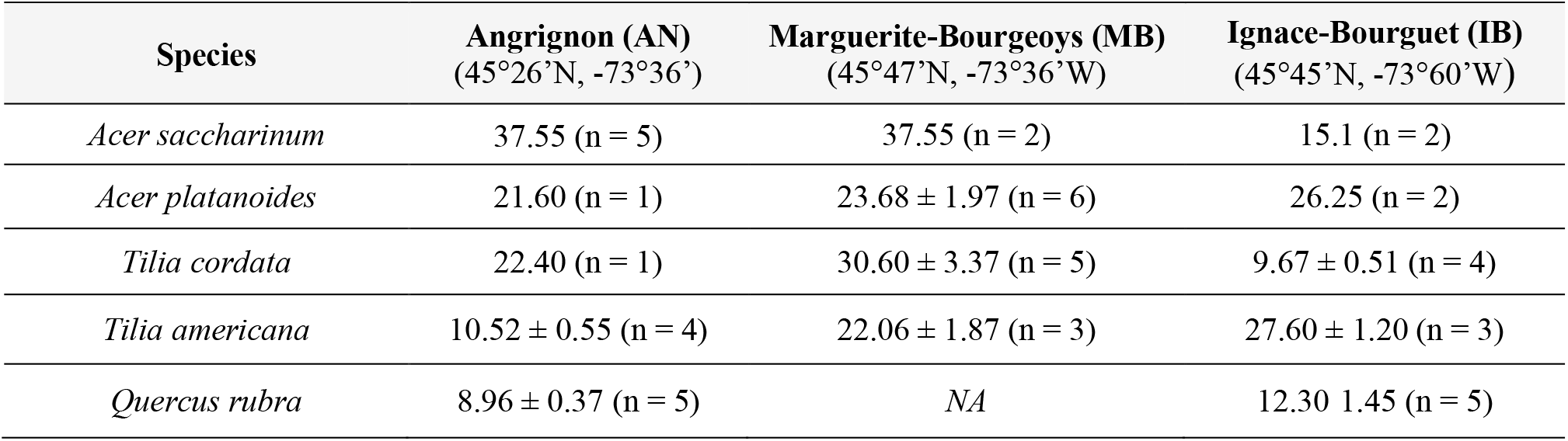
Mean (± SD) diameter at breast height (in cm) and number of trees initially selected for each park and species.

### Tree selection

Every tree in Angrignon, Ignace-Bourget and Marguerite-Bourgeoys parks had been previously geolocalized and identified to the species level. This information was accessible through the city database for urban trees (http://donnees.ville.montreal.qc.ca/dataset/arbres). We selected a total of 48 trees of five deciduous species (Table 1). Three species are native to the study area (*Acer saccharinum* L., *Tilia americana* L., *Quercus rubra* L.) while two are exotics, from Europe (*Acer platanoides* L., *Tilia cordata* Mill.). These species are amongst the most abundant tree species in the city of Montreal where together they represent 37% of all the tree species of the public domain. In agreement with the city of Montreal administration, we only selected trees with a diameter at breast height (DBH) greater than 8 cm (mean ± SD: 18.38 ± 9.36) (to withstand the sampling of leaves required for the experiment) and with low branches that could be easily accessed using a stepladder (for safety).

### Predation rate assessment

We used artificial caterpillars made with modelling clay to estimate predation rate on sampled trees (Ferrante, Lo Cacciato, & Lovei, 2014; Howe, Lövei, & Nachman, 2009). We installed 15 artificial caterpillars per tree. We haphazardly selected three low (2.5-3.5 m from ground) branches facing different directions and installed five artificial caterpillars per branch (total: 720 caterpillars). Caterpillars were 3 cm long, and modelled to match the approximate form and size of real caterpillars. They were modelled using a 1-cm ball of non-toxic and odourless green modelling clay *(Sculpey III String Bean colour)* and secured on thin branches using a 12-cm long, 0.5 mm diameter, non-shiny metallic wire.

We exposed artificial caterpillars for 11 days in late spring (from May 29^th^ to June 9^th^, 2018) and for 6 days in early summer (from July 18^th^ to July 24^th^, 2018). These seasons were chosen to cover the main activity period of both predators and herbivores. Artificial caterpillars were left untouched for the full duration of each survey. We estimated total predator attack rate as the number of artificial larvae with any predation mark, divided by the total length of the observation period in days. There were uncertainties regarding predator identity responsible for predation marks. Most of the marks were attributable to birds or arthropods, while very few were attributable to small mammals, therefore, we chose to combine predation marks primarily attributed to birds or arthropods into a single category, which we refer to as total predation.

Branches of three trees were accidentally pruned by city workers in late spring so that the predation rate could not be estimated on these trees for the first survey. Three new trees of the same species were selected for the second survey, in early summer.

### Leaf insect herbivory

We estimated insect herbivory on leaves (Kozlov et al., 2017) as the percentage of leaf area removed or impacted by insect herbivores (including chewing, skeletonizing and mining damage, collectively referred to as ‘herbivory’). At the end of the second predation survey, we collected 10 leaves per branch on the same branches on which we had exposed artificial caterpillars, starting with the most apical, fully-developed, leaf to the 10th leaf down to branch basis (Total: 30 leaves per tree). We estimated total herbivory (i.e., total leaf area consumed or impacted by herbivores, regardless of their identity) as well as damage made by chewing, mining and sap-feeding herbivores at the level of individual leaves by using an ordinal scale of eight percentage classes of defoliation: 0%; 0-1%, 1-5%; 6-10%; 11-25%; 26-50%; 51-75% and 76-100%. We counted the number of galls per leaf. Most damage was made by leaf chewers, while other damage had a skewed distribution, preventing detailed analyses for each type of damage separately. We therefore analysed total herbivory by averaging herbivory at the level of individual trees and using the median of each class of defoliation. Herbivory was scored by a single observer (BC), who was blind to tree identity.

### Tree neighbourhood

We used three variables to describe tree neighbourhood in a 20-m radius around each focal tree: tree density (defined as the number of neighbouring trees in that radius), tree species diversity (Shannon diversity index) and the number of conspecific trees around each focal tree. Those variables were obtained using QGIS Geographic Information System software (QGIS Development Team, 2018). Excluding focal tree species, the most common tree species in the vicinity of focal trees were the smooth serviceberry (*Amelanchier leavis* Wiegand), the white spruce (*Picea glauca* Voss), the green ash (*Fraxinus pennsylvanica* Marshall) and the eastern cottonwood (*Populus deltoides* Marshall), all of them native to the region. We should note that, as focal trees were not necessarily 20m or more apart, we could not avoid that some “neighbour” trees were used in more than one neighbourhood, and some focal trees were also within the neighbourhood of another focal tree.

### Statistical analyses

We used the information theory framework to identify the best model fitting our data and applied model averaging whenever necessary to estimate model coefficient parameters (Grueber, Nakagawa, Laws, & Jamieson, 2011). We first built a full model including tree density (*Density*), tree diversity (*Diversity*), number of conspecifics (*Conspecific*), origin of the focal tree (*Origin*, native of exotic), park (*Park*), and predation rate (*Predation*) as fixed factors and tree species identity (*Species*) as a random factor:

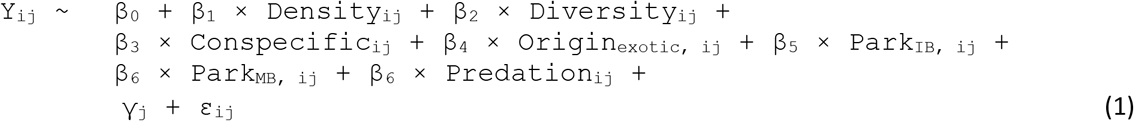

Where Y_ij_ is the herbivory on tree individual *i* in tree species *j, β* are model coefficient parameters for fixed effects, γ_j_ is the random effect of tree species identity and ε the residuals.

To ease the interpretation of parameter estimates after model averaging, we standardized the input variables using Gelman’s approach (Gelman, 2008). We then applied a procedure of model selection based on the Akaike’s criterion corrected for small sample size (AICc) by running every model nested within the full model. As tree density and tree diversity were correlated (Pearson’s correlation: *r* = 0.71), we excluded all sub-models that included these predictors together. We ranked all models based on difference in AICc between each model and the top ranked model with the lowest AICc (ΔAICc). Models with a ΔAICc < 2 are generally considered equally supported by the data or not differentiable from the top ranked model. Finally, we estimated model fit by calculating marginal (R^2^m) and conditional (R^2^c) R^2^ values, corresponding to variance explained by fixed effects only (R^2^m) and by fixed and random effects (R^2^c) (Nakagawa & Schielzeth, 2013). When multiple models had a ΔAICc < 2, we used a model averaging approach to build a consensus model including all variables found in the set of best models. We considered that a given predictor had a significant effect if its 95% confidence interval did not overlap zero. When only one model had a ΔAICc < 2, we used it as the best model. We used a square-root transformation of insect herbivory to satisfy model assumptions of normality and homogeneity of residuals.

We used the same approach to test the effect of tree neighbourhood on predation rate, log-transforming predation rate to satisfy model assumptions. Model equation (2) included the fixed effect of sampling season (*Season*) and the random effect of tree identity (τ_k_), nested within tree species identity as an additional random factor accounting for repeated measurements of the same individuals:

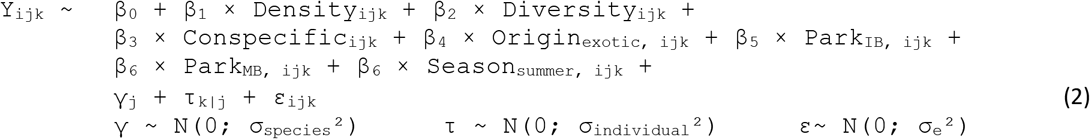

Statistical analyses were performed using the R software version 3.4.4 (R Core Team 2019) with packages *lme4* (Bates, Mächler, Bolker, & Walker, 2015) and *MuMIn* (Barton 2019).

## Results

### Insect herbivory

Herbivory was on average (± SE) 7.19 ± 0.70 % (*n* = 48). Leaf damage was lower in *Acer platanoides* (3.53 ± 0.54) and *A. saccharinum* (3.86 ± 0.47) than in *Quercus rubra* (8.77 ± 1.65), *Tilia americana* (10.3 ± 1.37) and *T. cordata* (8.75 ± 1.75) (Fig. 1A).

**Figure 1.**
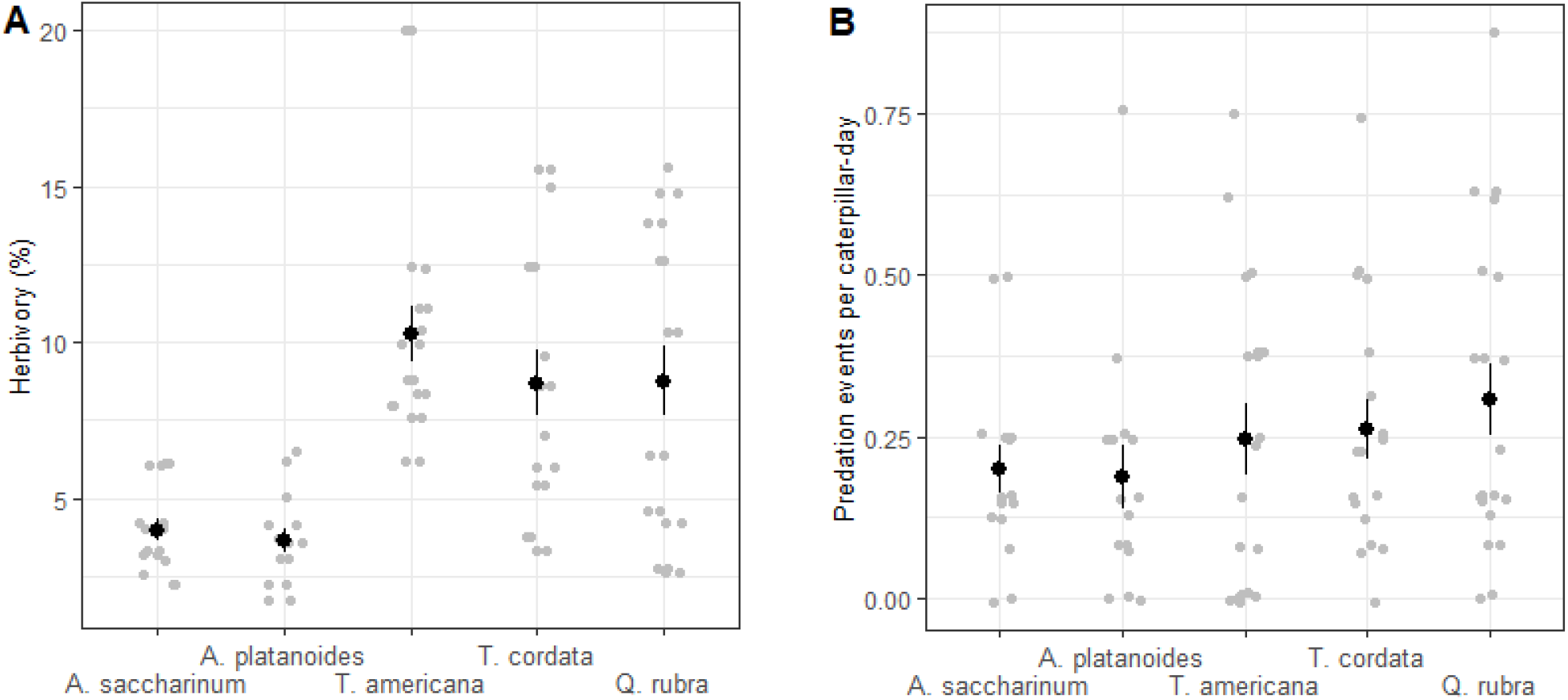
Effect of tree species identity on insect herbivory (A) and predation rate (B). Black dots and solid lines represents mean ± SE calculated on raw data. Herbivory is the percentage of leaf area removed or impacted by herbivores in early summer. Predation events per caterpillar-day is the number of caterpillars attacked per day in late spring

There were six models competing with the top ranked model in a range of 2 units of ΔAICc (Table 2). These models included tree Shannon diversity, predation rate and tree origin as predictors. Insect herbivory decreased significantly with increasing tree diversity (average model coefficient parameter estimate ± CI: – 0.482 ± [-0.91;-0.05], Fig. 2A, Table 3) and with increasing predation rate (–0.473 ± [-0.91; -0.003]) (Fig. 2B, Table 3). Among the set of best models, fixed effects explained between 7 and 12% of variability in insect herbivory. Fixed and random effects together explained between 47 and 65% of variability in insect herbivory.

**Table 2.** Summary results of model selection of tree neighbourhood effect on herbivory rate: set of models with ΔAICc < 2. Only predictors that were present at least once in the set of best models are represented. R^2^m and R^2^c represent fixed and fixed *plus* random factor, respectively.

| Model | Model covariates |  |  |  | Model selection |  |  |  |
| --- | --- | --- | --- | --- | --- | --- | --- | --- |
| | Intercept | Predation | Origin | Diversity | K | Log L | $\Delta AICc$ | $R^2m$ ($R^2c$) |
| **1** | 2.53 |  |  | -0.52 | 1 | -46.44 | 0.00 | 0.09 (0.46) |
| **2** | 2.52 | -0.52 |  | -0.44 | 2 | -45.18 | 0.04 | 0.12 (0.58) |
| **3** | 2.51 | -0.51 |  |  | 1 | -46.79 | 0.70 | 0.07 (0.56) |
| **4** | 2.53 | -0.44 | 0.171 | -0.43 | 3 | -44.64 | 1.67 | 0.12 (0.65) |
| **5** | 2.53 |  | 0.078 | -0.52 | 2 | -46.07 | 1.82 | 0.08 (0.53) |
| **6** | 2.53 | -0.53 | 0.357 |  | 2 | -46.12 | 1.92 | 0.08 (0.62) |

**Table 3.** Summary results after model averaging: effects of each parameter presents on the set of best models on herbivory rate. Bold parameter are significant. Relative importance is a measure of the prevalence of each parameter in each model used in model averaging.

| Parameter | Estimate | Adjusted SE | Confidence interval | Relative importance |
| --- | --- | --- | --- | --- |
| **(Intercept)** | 2.53 | 0.31 | (1.91, 3.14) |  |
| **Diversity** | -0.48 | 0.21 | (-0.91, -0.05) | 0.72 |
| **Predation** | -0.47 | 0.22 | (-0.91, -0.003) | 0.64 |
| Origin | 0.19 | 0.71 | (-1.20, 1.60) | 0.31 |

**Figure 2.**
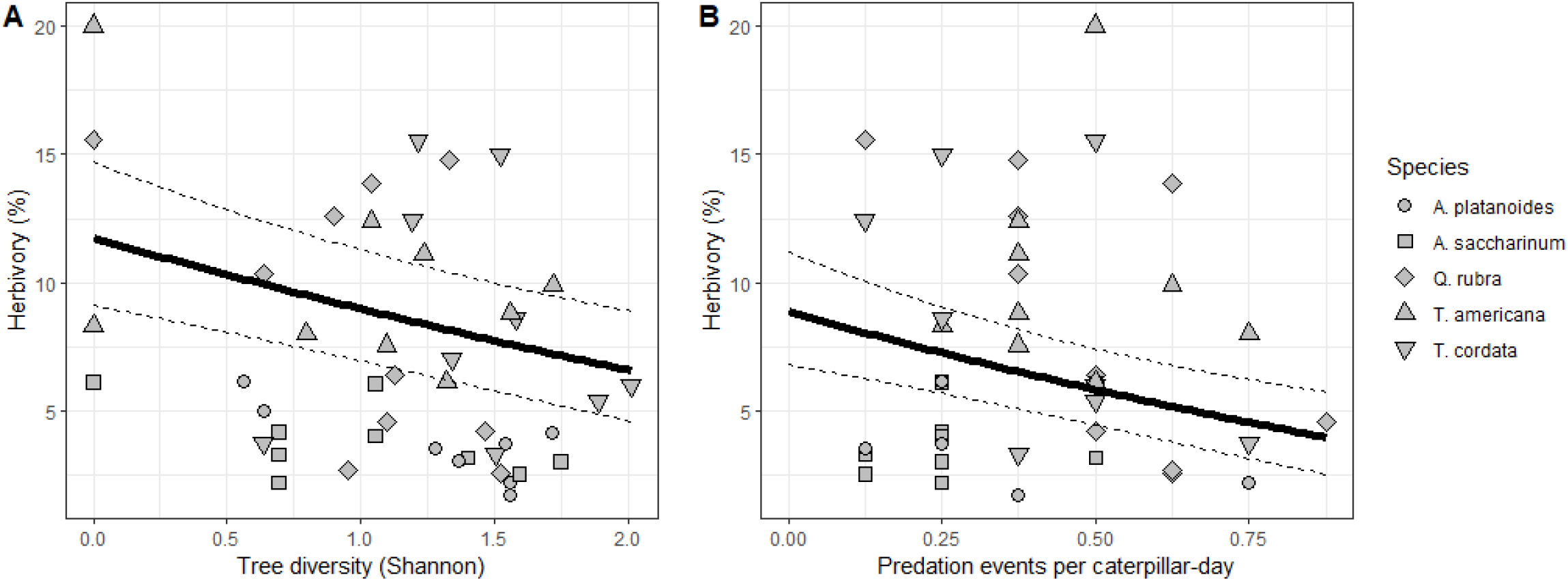
Effects of tree diversity (A) and predation rate (B) on insect herbivory. Solid and dashed lines represent prediction and adjusted standard error of the average model respectively (Table 3). Herbivory is the percentage of leaf area removed or impacted by herbivores in early summer. Tree diversity is represented by Shannon’s diversity index. Predation events per caterpillar-day is the number of caterpillars attacked per day in late spring.

### Predation

Of the 1,315 artificial caterpillars that we installed, 198 displayed marks unambiguously attributable to predators (*i.e*., 15%). Predation rate varied between 0 and 0.87 per caterpillar-day (Fig. 1B). Only one model had a ΔAICc < 2 and was thus selected as the best model. This best model included only Season, with predation rate two times higher in late spring (mean ± CI: 0.44 ± [0.31, 0.58] caterpillars·day^-1^) than in early summer (0.20 ± [0.16, 0.24] caterpillars·day^-1^). Season explained 56 % of variability in predation rate and, collectively, fixed and random effects explained 59 % of variability in predation rate.

## Discussion

We confirmed that tree diversity can influence insect herbivory on urban trees. Specifically, we found that insect herbivory decreased with increasing tree diversity providing support for the associational resistance hypothesis (Castagneyrol et al., 2017). We also found a negative correlation between predator attack rate and insect herbivory. Although further analyses are needed to confirm this relationship, our findings provide support for the view that increasing tree diversity can enhance regulation of insect herbivores by natural enemies in urban forests.

Our results are in line with several studies having reported reduced herbivory in trees surrounded by heterospecific neighbours (reviewed by Castagneyrol et al., 2014; Jactel et al., 2017). It also adds to the growing number of studies documenting diversity-resistance relationships in urban environments (Clem & Held 2018; Doherty, Meagher, & Dale 2019; Frank 2014). However, it conflicts with other results suggesting an increase in herbivore abundance with increasing plant diversity in urban environments (Mata et al., 2017), although the relationship between herbivore abundance and actual herbivory is not always positively correlated (Barbosa et al., 2009; Schueller, Paul, Payer, Schultze, & Vikas, 2019). Tree diversity may have influenced the probability of focal trees being found and colonized by herbivores. Theory predicts that specialist herbivores have greater difficulties finding their host trees when they are surrounded by heterospecific neighbours (Castagneyrol et al., 2014; H. Jactel, Brockerhoff, & Duelli, 2009). It is possible that non-host neighbours disrupted the physical and chemical cues used by insect herbivores to locate their hosts (Damien et al., 2016; H. Jactel et al., 2011; Zhang & Schlyter, 2004). However, and contrary to our expectations, we did not find any significant effect of conspecific tree density on insect herbivory, thus ruling out the resource concentration hypothesis in this particular case. However, because our study was observational, we could not separate the effect of conspecific neighbour density from heterospecific neighbour density. In the absence of data on the identity of herbivores responsible for herbivory, further speculation would be hazardous.

Insect herbivory varied across tree species but did not differ between native and non-native species, thus not providing support for predictions of the enemy release hypothesis (Cincotta, Adams, & Holzapfel, 2009; Meijer, Schilthuizen, Beukeboom, & Smit, 2016). One possible explanation for this result could be that native herbivores spilled over exotic tree species from neighbouring native tree species, as it was recorded in previous studies (Branco et al., 2015). This would have been facilitated by the fact that exotic tree species (from Europe) had congeneric species in Canada. Although we only surveyed a handful of native and exotic species, making any generalization hazardous, we can speculate on the lack of difference between native and non-native species. It is also important to note that a large part of the variability in leaf insect damage was attributable to the species on which leaf samples were collected. In particular, both *Acer platanoides* and *A. saccharinum* were far less damaged than *Tilia cordata, T. americana* and *Quercus rubra*. In a recent study in Michigan, Schueller et al., (2019) also reported greater insect herbivory (and herbivore diversity) on *Quercus* species as compared to *Acer* species, which is consistent with the view that plant species identity can drive arthropods community and abundance on forest host trees (Burghardt, Tallamy, & Gregory Shriver, 2009; Pearse & Hipp, 2009).

We found a significant negative correlation between predator attack rate and insect herbivory measured later in the season. This finding suggests a potential relationship between herbivory and predation in urban environments (Faeth, Warren, Shochat, & Marussich, 2005; Kozlov et al., 2017 but see Long & Frank, 2020).

However, we refrain from concluding that predation was the main driver of insect herbivory for several reasons. First, the effect size of the herbivory-predation relationship was small, as was model R^2^ (Table 3). Second, concerns remain about how well predation on artificial prey represents of actual predation (Lövei & Ferrante, 2017; Rößler, Pröhl, & Lötters, 2018). In particular, artificial caterpillars used to assess predation rate modelled lepidopteran-like leaf chewing caterpillars and thus, caution is needed when it comes to extrapolate predator attack rates to other herbivore feeding guilds. Third, we had no information on actual natural prey density in focal and neighbouring trees. Yet, prey availability may have influenced the functional response of bird insectivores (e.g. optimal foraging) such that we cannot exclude that herbivory actually drove predation rate instead of the other way around. Finally, the putative effect of predation on herbivory may be weak in respect to other factors acting directly upon herbivores in urban environments such as drought (Huberty & Denno, 2004; Mattson, 1980; Meineke & Frank, 2018), extreme heat (Dale & Frank, 2014; Meineke, Dunn, Sexton, & Frank, 2013) and pollution leading to altered foliage quality (Kozlov et al., 2017; Mattson, 1980; Moreira et al., 2019).

Contrary to the important effect of tree species identity on insect herbivory, tree species had no clear influence on predation rate on dummy caterpillars, which contradicts the view that tree species identity can modulate attack rates of caterpillars by birds (Mooney & Singer, 2012; Nell et al., 2018). Variation in predator density between plants is often related to an indirect effect of the plant on the density (Bailey et al. 2006) or quality (Brower et al. 1967, Clancy and Price, 1987) of their preys (herbivores). However, such effect of plant identity is not relevant when using dummy caterpillars, as neither their abundance nor their quality can be affected by plant species identity, which could explain the contradiction between past results and our study.

Predation was greater during the first survey, in late spring, than during the second survey, in early summer. This result could be explained either by a lower foliage density in trees in spring, making it easier for predators to detect artificial caterpillars, or by greater predator activity matching the phenology of wild caterpillars and feeding period of chicks (Coley, 1980; Raupp & Denno, 1983). We cannot either exclude that birds learned to avoid artificial caterpillars, thus resulting in much lower predation pressure during the second survey. However, unless bird ability to avoid artificial caterpillar varied between tree species and neighbourhood, we do not see this possibility as a major threat to our inferences.

## Conclusion

Our study suggests several ecological factors drive leaf insect herbivory in the urban trees of the Montreal city. In particular, we found that insect herbivory decreased with both increasing tree diversity and predator activity. While biological invasions and global warming are increasing risks to urban trees, more and more cities choose to ban or reduce the use of pesticides in urban parks and green areas (Sustainable Use of Pesticides Directive 2009), such as in Montreal. In this context, diversifying urban tree cover in urban parks might help to reduce insect damage, which could result in a better provision of services provided by trees in cities (Beyer et al., 2014; Bowler, Buyung-Ali, Knight, & Pullin, 2010; Nowak, Hirabayashi, Bodine, & Greenfield, 2014).

## Data accessibility

Data, script, codes and supplementary material are available online from the Data INRAE repository: https://doi.org/10.15454/R4NESA

## Acknowledgements

This study has been carried out with financial support from (i) the French National Research Agency (ANR) in the frame of the Investments for the future Programme to BC, within the Cluster of Excellence COTE (ANR-10-LABX-45), (ii) the Conseil Franco-Québécois de Coopération Universitaire (FRQNT – Samuel-de-Champlain fund) to BC and AP, and (iii) a Discovery grant to AP from the Natural Sciences and Engineering Research Council of Canada. We thank Charles Desroches, Elyssa Cameron, Christian Messier, summer interns at UQAM, and the city of Montréal, Arrondissement du Sud-Ouest for their help with field work. We also thank Frédéric Barraquand and Benjamin Brachi for their constructive comments on the preliminary version of the paper. We thank Luc Barbaro and Steve Frank for their friendly reviews of early drafts of this paper. Finally, we are grateful to Ian Pearse and Freerk Molleman for their reviews of prior manuscript versions. Version 5 of this preprint has been peer-reviewed and recommended by Peer Community In Ecology (https://doi.org/10.24072/pci.ecology.100061)

